# Sexual dimorphism and the effect of wild introgressions on recombination in *Manihot esculenta*

**DOI:** 10.1101/794339

**Authors:** Ariel W. Chan, Amy L. Williams, Jean-Luc Jannink

## Abstract

Recombination has essential functions in evolution, meiosis, and breeding. Here, we use the multi-generational pedigree, consisting of 7,165 informative meioses (3,679 female; 3,486 male), and genotyping-by-sequencing (GBS) data from the International Institute of Tropical Agriculture (IITA) to study recombination in cassava (*Manihot esculenta*). We detected recombination events using SHAPEIT2 and duoHMM, examined the recombination landscape across the 18 chromosomes of cassava and in regions with known introgressed segments from cassava’s wild relative *Manihot glaziovii*, constructed a genetic map and compared it to an existing map constructed by the International Cassava Genetic Map Consortium (ICGMC), and inspected patterns of recombination placement in male and female meioses to see if there is evidence of sexual dimorphism in crossover distribution and frequency. We found that the placement of crossovers along chromosomes did not vary between the two sexes but that females undergo more meiotic recombination than males. We also observed that introgressions from *M. glaziovii* decreased recombination in the introgressed region and, in the case of chromosome 4, along the entire length of the chromosome that the introgression is on. We observed a dosage effect on chromosome 1, possibly suggesting the presence of a variant on the *M. glaziovii* haplotype that leads to lower overall recombination in the introgressed region.

## INTRODUCTION

Recombination plays essential roles in evolution, meioses, and breeding. Recombination generates genomic diversity in a population by creating new combinations of already existing alleles. In the context of meiosis, it aids in homology recognition and ensures proper segregation of homologous chromosomes [1]. Improper disjunction, or nondisjunction, results in aneuploidy, a deleterious outcome in which gametes have less or more than the typical chromosome number. Recombination also serves as an important breeding tool as its rate dictates the resolving power of quantitative trait mapping and the precision of allele introgression.

The number of crossovers per chromosome and the distribution of crossovers along chromosomes are tightly controlled. Crossover number appears to be constrained by both an upper and lower bound. The reason for a lower bound on crossover number is clear since in most species, there is a need for one obligatory crossover per tetrad to prevent aneuploidy [2]. The reason(s) for an upper bound on crossover number, however, is less obvious. Results from a recent study, where crossover rate was significantly increased in mutant *Arabidopsis thaliana*, suggest that reduced fertility (in the form of reduced pollen viability and seed set) may be associated with increased recombination [3]. One plausible evolutionary explanation for the existence of an upper bound on recombination is that beneficial alleles residing on the same haplotype may collectively act to increase fitness. Recombination can break these associations, resulting in reduced progeny fitness [4].

The distribution of crossovers along chromosomes is not random and is influenced by chromosome features such as chromatin structure, gene density, and nucleotide composition [5]. The occurrence of a crossover at one location also reduces the likelihood that another crossover will occur in close proximity. This nonrandom placement of crossovers, known as crossover interference, results in a pattern where recombination events appear more evenly spaced [6], [7]. Interference may therefore serve as a biological mechanism to ensure that every pair of homologous chromosomes undergoes at least one crossover event, which is necessary for proper disjunction.

In many species, crossover frequency and distribution along chromosomes differs between female and male meiosis, a phenomenon referred to as heterochiasmy [8]. The direction and degree of these differences are species-specific, and most extreme are cases in which one of the two sexes lacks meiotic recombination entirely. Male *Drosophila melanogaster*, for example, do not recombine during meiosis. To date, no investigation of sexual dimorphism has been conducted in cassava.

Cassava is a diploid organism with an estimated genome size of approximately 772 Mb spread across 18 chromosomes with the reference genome spanning 582.28 Mb [9]. The International Cassava Genetic Map Consortium (ICGMC) generated a consensus genetic map of cassava that combines 10 mapping populations [10]. The 10 mapping populations consisted of one self-pollinated cross and nine biparental crosses (14 parents total; 3,480 meioses). The genetic map is 2,412 cM in length and organizes 22,403 GBS markers on 18 chromosomes. Here, we used the multi-generational pedigree from the International Institute of Tropical Agriculture (IITA) to characterize recombination in cassava. We used duoHMM-corrected, SHAPEIT2-inferred haplotypes to detect SNP intervals flanking a crossover event then used these intervals to map the recombination landscape across cassava’s 18 chromosomes [11]. We built a genetic map from 7,165 meioses, compared it to ICGMC’s composite map, and constructed sex-specific genetic maps to see if crossover distribution and frequency differ significantly between the two sexes. To evaluate the impact of known genomic introgressions from *M. glaziovii* into *M. esculenta*, we also constructed introgression dosage-specific genetic maps. Finally, we examined if there is evidence of chromosomal interference.

## MATERIALS AND METHODS

### The IITA germplasm population structure

The IITA pedigree consists of 7,432 unique individuals. Each individual belongs to one of four breeding groups: Genetic Gain (GG; *n* = 494), TMS13 (*n* = 2,334), TMS14 (*n* = 2,515), or TMS15 (*n* = 2,089). Of the 494 GG individuals, 258 are the progeny of GG-GG crosses and the remaining 236 individuals are founders (individuals for which we do not have parent data). All TMS13 members are the progeny of GG-GG crosses. Of the 2,515 TMS14 individuals, 1,881 are the progeny of TMS13-TMS13 crosses and the remaining are GG-GG progeny. The TMS15 group consists of 920 progeny of TMS14-TMS14, seven of TMS13-TMS14, 1,159 of TMS13-TMS13, and three individuals of TMS13-GG crosses, respectively. All but the GG group were generated by the “Next Generation Cassava Breeding Project” (nextgencassava.org), hereafter referred to as “NextGen”.

### Merging replicate GBS records of each proband

We found GBS data for 7,294 of the 7,432 IITA individuals (*n*_*GG*_ = 366, *n*_*TMS13*_ = 2330, *n*_*TMS14*_ = 2509, and *n*_*TMS15*_ = 2089). Of the 366 GG individuals, 189 had more than one GBS record (i.e., NextGen sequenced these 189 individuals multiple times). Before merging the data from replicate sequence runs of an individual, we verified that no erroneous samples existed among the putative replicates (i.e. verified that all putative replicates derived from an identical individual) using BIGRED [12]. Using Bayes Theorem, BIGRED calculates the posterior probability distribution over the set of relations (i.e., source vectors) describing the putative replicates of an individual, and infers which of the samples originated from an identical genotypic source. Of the 189 GG BIGRED runs, 21 produced ambiguous results. An ambiguous BIGRED result occurs when BIGRED returns a source vector where no source has a clear majority. Because we were unable to resolve these cases, we excluded these 21 GG individuals from future analyses. We merged the allelic depth data for the 168 GG individuals with unambiguous BIGRED results, merging only the samples that were inferred to be true replicates. We repeated this process for TMS13 and TMS14 individuals (all individuals in the TMS15 group were sequenced once). Of the 2,330 TMS13 individuals, 156 had more than one GBS record and 10 produced ambiguous BIGRED results. Of the 2,509 TMS14 individuals, 62 had more than one GBS record, and three produced ambiguous results. We excluded these 13 TMS13 and TMS14 individuals from further analyses. Table 1 summarizes these results.

**Table 1.** Summary of data records for each breeding group. The table shows the number of individuals, the number of individuals that were sequenced more than once, and the number of individuals with ambiguous BIGRED results for each breeding group.

| Group ID | Number of individuals in group | Number of individuals with >1 sequence record | Number of individuals with ambiguous BIGRED results |
| --- | --- | --- | --- |
| GG | 366 | 189 | 21 |
| TMS13 | 2330 | 156 | 10 |
| TMS14 | 2509 | 62 | 3 |
| TMS15 | 2089 | 0 | 0 |

### Validating IITA pedigree records using AlphaAssign

Of the remaining 345 (=366-21) GG individuals listed in the pedigree, 187 GG individuals had at least one listed parent with available GBS data. These parents also belong to the GG population. We used the parentage assignment algorithm AlphaAssign to validate the existing pedigree information for these 187 GG individuals [13]. AlphaAssign frames the parentage assignment problem as a relationship classification problem. Rather than directly attempting to identify target individual *t*’s parent from a list of candidate individuals, AlphaAssign attempts to classify the relationship between target individual *t* and each candidate individual *c*. AlphaAssign considers four possible target-candidate relationships, *H*: the candidate individual is (1) a parent of the target individual, (2) a full-sibling of the target individual’s parent, i.e., the target individual’s uncle, (3) a half-sibling of the target individual’s parent, and (4) unrelated to the target individual. Specifically, AlphaAssign calculates the posterior probability of these four relations, given the observed allelic depth data of *c* and *t* (and if known, the allelic depth data for a known parent of *t*; if individual *t* has no known parent, the algorithm makes use of a ‘dummy parent’ whose genotype probabilities at a given site are calculated using estimated allele frequencies and assuming Hardy-Weinberg Equilibrium).

Because AlphaAssign looks at the relationship between pairs of individuals rather than among triplets, we needed to run AlphaAssign two times to validate IITA’s pedigree information. We walk through the validation procedure for GG individuals. In the first run, we provided the algorithm no pedigree information (i.e., all calculations involved the use of a dummy parent). For each target individual, we listed all GG individuals as candidate parents (we did not list an individual as its own candidate parent). We fed the algorithm allelic depth data from 1,000 randomly sampled sites across cassava’s 18 chromosomes. To filter for LD, we sampled sites such that no two sites fell within 20 kb from one another. For each target individual, we identified the candidate individual with the highest score statistic and listed this top-scoring candidate as the target individual’s parent in a (newly created) pedigree file. We ran AlphaAssign a second time, this time providing AlphaAssign with pedigree information, i.e., the AlphaAssign-inferred pedigree generated from the results of the first run. We again identified the candidate individual with the highest score statistic for each target individual. Upon completing the two runs, each target individual had two AlphaAssign-inferred parents. We compared the AlphaAssign-inferred pedigree with IITA’s existing pedigree. We repeated this analysis for the TMS13, TMS14, and TMS15 group and present the results of all four breeding groups in Table 2. We built the list of candidate individuals for each breeding group based on how IITA generated each group (see above). For TMS13 target individuals, we listed all GG individuals as candidate parents. For TMS14 target individuals, we listed all GG and TMS13 individuals as candidate parents. For TMS15 target individuals, we listed all GG, TMS13, and TMS14 individuals as candidate parents.

**Table 2.**
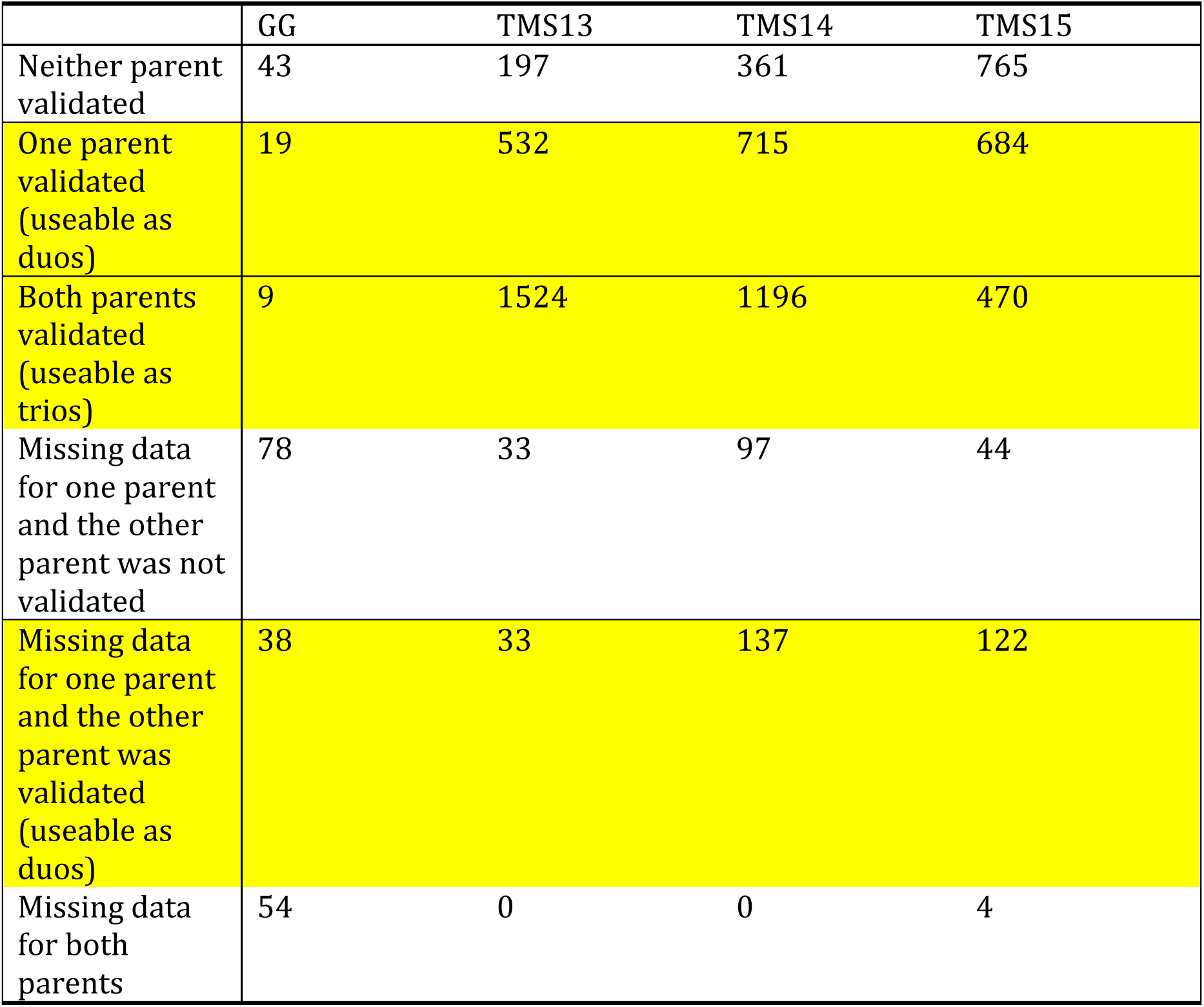
Pedigree validation results from AlphaAssign. The table shows the results of our pedigree validation procedure. Rows highlighted in yellow represent useable data (either duos or trios). An individual’s data is labeled “missing” when we either could not find GBS data for that individual or when we could not resolve their BIGRED results.

### Filtering the GBS allele depth data before calling genotypes

We have allelic depth data for each individual at each site. The allelic depth data for individual *d* at site *v* is a record of the observed counts of each of the two alleles in individual *d* at site *v*: 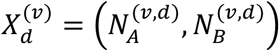, where 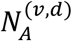 and 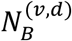 denote the observed counts of allele A and allele B, respectively, in individual *d* at site *v*. We removed sites with >70% missing data then calculated the proportion of missing data for each individual and removed individuals with >80% missing data. Here, we defined “missing” as observing zero reads for a given individual at a given site. The filter removed one individual IITA-TMS-IBA011610 from analysis. Exclusion of this individual causes offspring IITA-TMS-IBA062021 to have no listed father or mother. We included IITA-TMS-IBA062021 in the analysis when phasing and imputing. Removal of this duo is inconsequential since this duo provides an uninformative meiosis (see the section “*Filtering the SHAPEIT2-duoHMM output*” below for a discussion of informative meioses). We then removed sites with a mean depth (calculated across all samples) greater than 120 to avoid spurious genotype calls within repeat regions, i.e., paralogs.

### Generating input data files for SHAPEIT2

SHAPEIT2 takes called genotypes as input. To obtain a set of called genotypes for our sample, we first calculated genotype posterior probabilities for each individual at each site. Given observed data 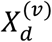 and fixed sequencing error rate *e* = 0.01, we computed the likelihood for genotype 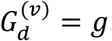. We calculated genotype likelihoods for a single individual at a single site, independent of all other individuals and sites in the sample, using the following equation:

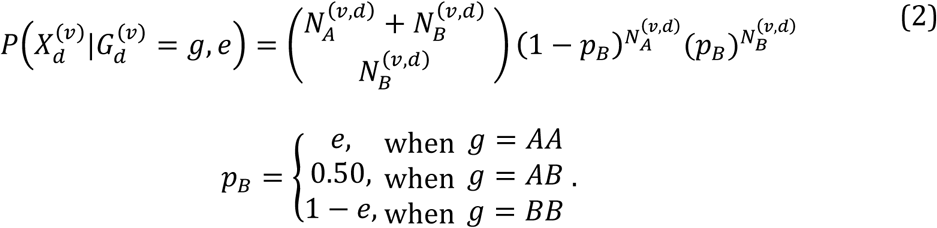

We estimated posterior probabilities for the three genotypes using the likelihoods defined above and assuming a genotype prior (then normalizing). This genotype prior varied depending on whether individual *d* had zero validated parents (i.e., was a founder), had one validated parent, or had two validated parents. If individual *d* had zero validated parents, we calculated its genotype prior for site *v* using the estimated frequency of the reference allele at site *v* and assuming Hardy-Weinberg Equilibrium (HWE). If individual *d* had one validated parent, we calculated its genotype prior for site *v* using the posterior probability distribution of its known parent, the genotype probability distribution of a ‘dummy’ parent, and the rules of Mendelian inheritance. We calculated the genotype probability distribution of the dummy parent at site *v* by using the estimated allele frequency at site *v* and assuming HWE. If individual *d* had two validated parents, we used the posterior probability distributions of its known parents and Mendelian inheritance rules to calculate individual *d*’s prior. Notice that this scheme requires calculation of posterior genotype probabilities in a sequential manner, propagating information down the pedigree to subsequent generations.

We called genotypes from these estimated posterior genotype probabilities, calling a genotype for individual *d* at site *v* only if one of the three possible genotypes had a posterior probability greater than or equal to 0.99. To qualitatively examine how SHAPEIT2 performs at different levels of missing data, we generated two datasets: one dataset where we removed sites that had more than 20% missing data and another where we removed sites that had more than 30% missing data We observed that when more markers are retained, SHAPEIT2-duoHMM detected a larger number of crossovers but crossover intervals were longer (a recombination event can only be resolved down to the region between its two flanking heterozygous markers in the parent). Results from the 20% dataset were very noisy, so we selected the 30% dataset to analyze. Table 3 shows the number of sites after applying the 30% maximum-missing filter for each chromosome. Supplementary Figure 1 shows the plots for each chromosome’s duoHMM-inferred crossover intervals for the 20% and 30% maximum-missing datasets.

**Table 3.** Number of sites remaining after filtering. The table lists the number of sites in the dataset before and after application of the 30% maximum-missing filter for each chromosome and removing monomorphic and singleton sites.

| Chromosome | Number of sites before filter | Number of sites after 30% maximum-missing filter | Number of sites after removing monomorphic and singleton sites |
| --- | --- | --- | --- |
| 1 | 30575 | 17159 | 3739 |
| 2 | 24151 | 14818 | 2265 |
| 3 | 22156 | 14493 | 2458 |
| 4 | 18271 | 9980 | 2519 |
| 5 | 20750 | 14106 | 1681 |
| 6 | 20174 | 12907 | 1609 |
| 7 | 12226 | 7163 | 1114 |
| 8 | 17991 | 11549 | 1689 |
| 9 | 18267 | 12194 | 1341 |
| 10 | 14985 | 8206 | 1573 |
| 11 | 18816 | 11267 | 1777 |
| 12 | 15906 | 9921 | 1703 |
| 13 | 15851 | 9969 | 2018 |
| 14 | 20635 | 11957 | 2939 |
| 15 | 20048 | 13239 | 1705 |
| 16 | 15447 | 9832 | 1267 |
| 17 | 15390 | 8716 | 2206 |
| 18 | 15053 | 9063 | 1524 |

**Figure 1.**
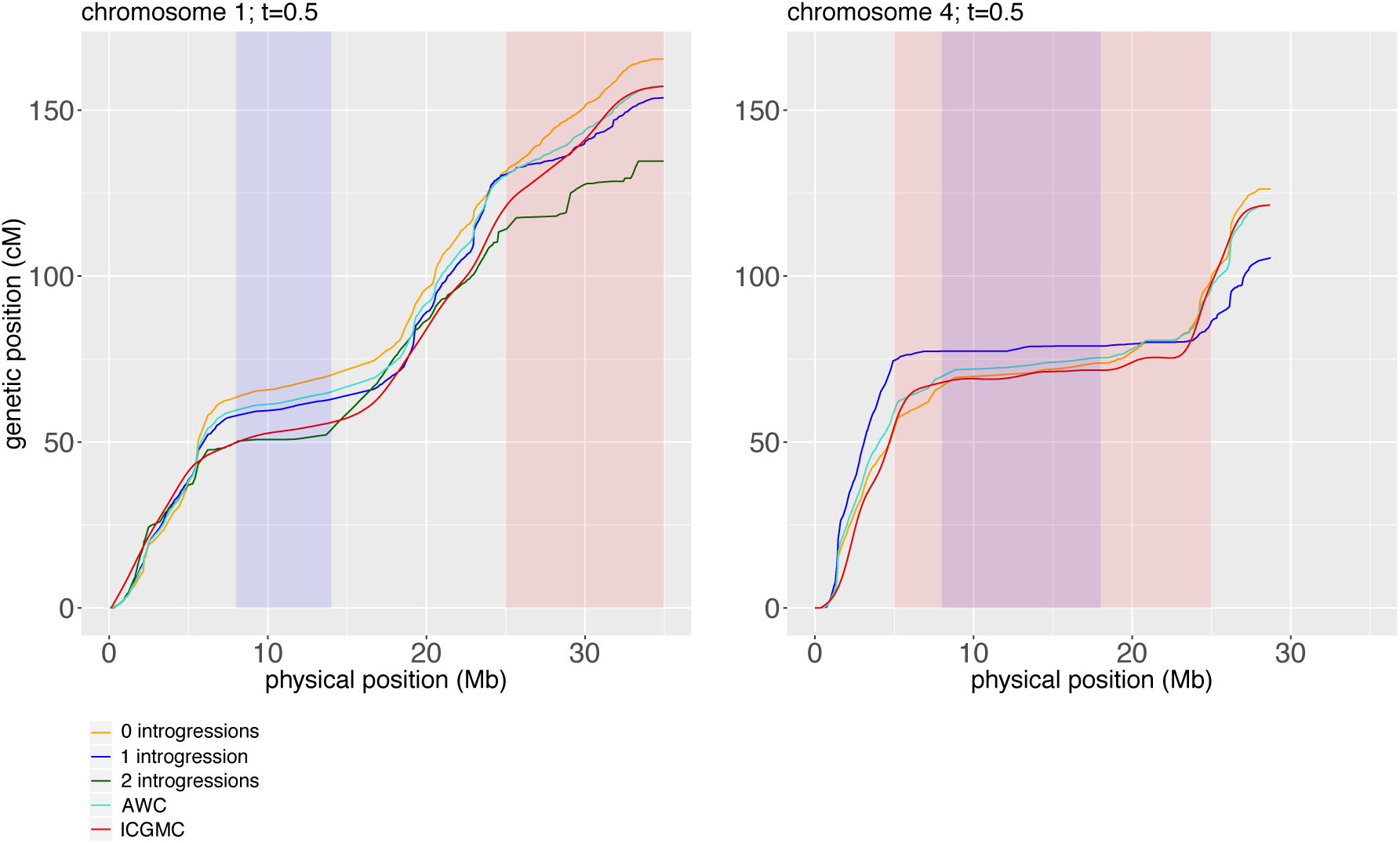
Comparison of our constructed genetic maps with the ICGMC map. We plotted the genetic position of our GBS markers and ICGMC’s markers as a function of physical position (Mb) for chromosomes 1 and 4. The left plot shows the maps for chromosome 1 and the right for chromosome 4. Centromeric regions of chromosomes are shaded in purple, and introgressed regions are shaded in red. We show five maps: ICGMC’s map (red), our sex-averaged genetic map constructed from the crossovers detected in all informative parents (turquoise; labeled AWC), and three genetic maps constructed from the crossovers detected in informative parents homozygous non-introgressed (orange), heterozygous for an introgression (blue), and homozygous introgressed (green).

### Running SHAPEIT2 and duoHMM

We used SHAPEIT2 and duoHMM to detect SNP intervals flanking a crossover event (a recombination event can only be resolved down to the region between its two flanking heterozygous markers in the parent). We followed the recommendations of O’Connell et al. [11] to phase and impute individuals in our pedigreed population. We ran SHAPEIT2, ignoring all explicit family information then applied duoHMM to combine the SHAPEIT2-inferred haplotypes with the verified family information to correct switch errors (SEs). The duoHMM Hidden Markov Model (HMM) was developed to detect recombination events and correct SEs in SHAPEIT2-inferred haplotypes. The true pattern of gene flow at each site is unobserved, and duoHMM infers the true inheritance states from the (imperfect) observed parental and child haplotypes. We refer the reader to the duoHMM paper for full specification of the HMM [11]. After estimating parameters of the HMM using the Forward Backward algorithm, duoHMM finds the most likely state sequence using the Viterbi algorithm. When duoHMM infers a SE in the Viterbi sequence in either the parent or child, duoHMM corrects the haplotypes by switching the phase of all loci following the SE. The algorithm applies these corrections sequentially down through each pedigree. We carried out both steps internally within SHAPEIT2 by using the ‘—duohmm’ flag.

To run SHAPEIT2 with the ‘duoHMM’ flag, we provided the algorithm with a set of genotypes, a genetic map, and verified pedigree information. SHAPEIT2 outputs either a single set of estimated (most-likely) haplotypes or a haplotype graph that encapsulates the uncertainty about the underlying haplotypes. We chose the latter output. SHAPEIT2 has multi-threading capabilities, but we chose not to use this feature in order to maximize the number of individuals that SHAPEIT2 conditions on during Gibbs sampling. We ran SHAPEIT2 with 14 burn-in iterations, 16 pruning iterations, and 40 main iterations. We increased the number of conditioning states to 200 states per SNP. The developers found it slightly advantageous to use a window size larger than 2 Mb when large amounts of identical by descent (IBD) sharing are present. We used a window size of 5 Mb. We provided SHAPEIT2 a genetic map that specifies the recombination rate between SNPs. We generated this genetic map by interpolating genetic distances of GBS markers using ICGMC’s composite genetic map. We used the default value of 15,000 for the effective population size, a parameter that scales the recombination rates that SHAPEIT2 uses to model patterns of LD.

### Detecting recombination events using duoHMM

Once duoHMM corrected SEs in the SHAPEIT2-inferred haplotypes, we reran duoHMM to infer recombination events. The HMM infers recombination events by calculating the probability of a recombination event between markers [11]. To detect crossovers, we sampled a haplotype pair for each individual from SHAPEIT2’s diploid graph then calculated the probability of a recombination event between pairs of markers. We repeated this process a total of 10 times then averaged the inter-SNP recombination probabilities across the 10 iterations. We included a crossover interval in subsequent analyses if the interval had a probability greater than or equal to *t* = 0.5. Supplementary Figure 1 shows those crossover intervals with probabilities greater than or equal to *t* = 0.9.

### Filtering the SHAPEIT2-duoHMM output

The power to detect recombination events is dependent on the structure of the pedigree. In a nuclear family with >2 offspring, most crossover events should be detectable, and we classify these pedigrees as *informative* towards recombination. We analyzed data from only those pedigrees having “informative” meioses, which we defined as a nuclear family consisting of >2 offspring or a pedigree consisting of three generations. We refer to the parents of these pedigrees as “informative parents” and the meioses in these pedigrees as “informative meioses”. Of the total 8,678 meioses in the data set, 7,165 were informative (3,679 female meioses; 3,486 male meioses).

### Building sex-averaged genetic maps

To build a genetic map for each chromosome, we first calculated the number of crossover events that occurred between each pair of SNPs. If a crossover event spanned multiple SNP intervals, we assigned a fraction of the crossover event to each of the spanned intervals, calculated as 1/(length of the SNP interval in bps). We then calculated the genetic length of each SNP interval on chromosome *y* by dividing the number of crossovers in each interval by a scaling factor *n*_*y*_, where *n*_*y*_ = (the genetic length of chromosome *y* in the ICGMC map)/(the total number of crossovers we detected on chromosome *y*). We did this so that our genetic map length of each chromosome is the same as ICGMC’s.

### Examining evidence of sexual dimorphism

We next examined the distribution of crossover events along each chromosome for female and male meioses separately. We divided each chromosome into windows of 1-Mb and determined the number of male meiotic crossovers and <u>female</u> meiotic crossovers in each window. To examine if crossover counts in each window varied between the sexes, we performed a chi-square test of equal counts in each window. To calculate the expected number of male crossovers in a given window, we calculated the proportion of total meioses analyzed that were male (i.e., 3,486/(3,679 + 3,486)) then multiplied this value by the total number of crossovers in the window. We calculated the expected number of female crossovers in a given window in the same way. We did not test for statistical significance in the last window of any chromosome since the last window is shorter than 1-Mb (no chromosome is perfectly divisible by 1-Mb). We could not perform the chi-square test for four of the 510 windows because these windows had one or more classes with an expected frequency count of less than five. We tested each window at a Bonferroni-corrected significance level of *α*/*m*, where *α* = 0.05 and *m* = 506 (i.e., the total number of windows tested). We also performed this test genome-wide at a significance level of 0.05.

### Examining if crossover placements are random and independent events

If crossover placements are random and independent events, the distribution of the number of crossovers observed on a given chromosome in a given parent-offspring pair is expected to follow a Poisson distribution. We used the deviance goodness of fit test to determine if crossover placements are Poisson distributed. For each chromosome, we performed a Poisson regression where we modeled the number of crossovers observed in a given parent-offspring pair *Y* as a function of the covariates “parent” and “sex”. The “parent” covariate specifies the parent involved in the parent-offspring pair, and the “sex” covariate specifies whether the parent was a female or male (i.e., were the crossovers observed in a male or female meiosis). We used the residual deviance to perform a chi-square goodness of fit test for the overall model. The residual deviance is the difference between the deviance of the current model and the maximum deviance of the ideal model where the predicted values are identical to the observed. If the residual difference is small enough, the goodness of fit test will not be significant, indicating that the Poisson model fits the data. We performed these test at a Bonferroni-corrected significance level of *α*/*m*, where *α* = 0.05 and *m* = 18 (i.e., the total number of chromosomes tested).

### Examining recombination patterns in introgressed regions on chromosome 1 and 4

Wolfe et al. detected large introgressed *M. glaziovii* segments on chromosome 1 and 4 in a collection of modern cassava landraces and elite germplasm [14]. They detected the largest introgressions on chromosome 1, spanning from 25 Mb to the end of the chromosome, and on chromosome 4 from 5 Mb to 25 Mb. To examine the recombination patterns in the introgression region on chromosome 1, we categorized our informative parents into three groups: individuals that are homozygous non-introgressed, heterozygous for introgressions, and homozygous introgressed. We obtained the introgression data for our samples from Marnin et al. All individuals listed as a parent in our verified pedigree were individuals analyzed in the Marnin et al. study, except individual TMS13F1079P0007, so we excluded TMS13F1079P0007 from our analysis. We briefly describe the data but refer the reader to the Marnin et al. manuscript for complete details. Marnin et al. defined a set of Introgression Diagnostic Markers (IDMs) across the cassava genome by comparing a panel of pure, non-admixed *M. glaziovii* individuals with a panel of pure, non-admixed *M. esculenta* individuals. A SNP belongs to the set of IDMs if either (1) the SNP is fixed for different alleles between the *M. glaziovii* and *M. esculenta* reference panels or (2) the SNP is fixed among *M. esculenta* samples but polymorphic in the *M. glaziovii* sample. Marnin et al. determined the number of *M. glaziovii* alleles each individual had at each IDM then divided each individual’s chromosomes into non-overlapping windows of 250-kb. For each individual, Marnin et al. determined the mean *M. glaziovii* dosage at each window by averaging across the IDMs falling within each window for that individual. To classify an individual as homozygous non-introgressed, heterozygous for introgressions, or homozygous introgressed on chromosome 1, we found the windows spanning the introgression region (25 Mb to 35 Mb) and calculated the mean *M. glaziovii* dosage in that region. Values ranged from 0 to 2, inclusive. We rounded dosages falling in the range [0,0.5), [0.5,1.5), and [1.5,2] to, respectively, 0, 1 and 2 *M. glaziovii* introgressions. We determined the number of crossover intervals overlapping the introgression region and performed a chi-square test of equal counts to examine if individuals with different numbers of introgressions experience different levels of recombination in the region at a Bonferroni-corrected significance level of 0.05/2. To calculate the expected number of crossovers for individuals that are homozygous non-introgressed, we calculated the proportion of informative meioses contributed by non-introgressed individuals (i.e., 2645/(2645+4047+439)) then multiplied this value by the total number of crossovers found in the introgressed region across all meioses. We calculated the expected number of crossovers for individuals heterozygous and homozygous for introgressions in the same way. We repeated this analysis for the introgression region on chromosome 4. We found no individuals that were homozygous introgressed on chromosome 4. To see if introgression status affected recombination frequency across regions of the chromosome with no introgression, we performed this analysis for the non-introgressed portion on chromosome 1 and 4, again using a significance threshold of 0.025.

### Building introgression-specific genetic maps

To examine how the number of introgressed segments from *M. Glaziovii* affected recombination on chromosomes 1 and 4, we constructed three additional genetic maps for each of the two chromosomes: one map constructed using the crossovers detected in individuals that are homozygous non-introgressed (0 introgressions), one constructed using individuals heterozygous for introgressions (1 introgression), and one constructed using individuals that are homozygous introgressed (2 introgressions). We followed the same procedure as before to build these maps (refer to the section “*Building sex-averaged genetic maps*”) but scaled the 0, 1, and 2 introgression maps such that their weighted average equaled the sex-averaged map. We walk through the scaling procedure for the 0 introgression map for chromosome 1. To calculate the genetic length of each SNP interval on chromosome 1’s 0 introgression map, we divided the number of crossovers (detected in homozygous non-introgressed parents) in each interval by the same scaling factor as before *n*1 (refer to the section “*Building sex-averaged genetic maps*” for a description of *n*1) but then multiplied this value by *m*, where *m* = (the total number of informative meioses used to construct the 0, 1, and 2 introgression maps)/(the number of informative meioses used to construct the 0 introgression map).

## DATA AVAILABILITY

Raw data and results are publically available on the Cassavabase FTP: ftp://ftp.cassavabase.org/manuscripts/Chan_et_al_2019/. The README file provides a description of each file. Supplementary Figures are available at FigShare.

## RESULTS

Using SHAPEIT2 and duoHMM, we detected a total of 67,833 and 51,741 crossover-containing intervals from female and male meioses, respectively, across the 18 chromosomes. Using these crossover intervals, we constructed a sex-averaged genetic map, which we compared to an existing map constructed by ICGMC. Our sex-averaged map has a median resolution of 420,366 bp.

To compare our map to ICGMC’s, we plotted the genetic position (cM) of our markers and ICGMC’s markers as a function of physical position (Mb). Figure 1 shows the maps for chromosomes 1 and 4. We show the plots for each chromosome in Supplementary Figure 2. At the qualitative level, the distribution of crossovers observed in our map is in good agreement with the ICGMC map. We found that crossovers are suppressed around centromeric regions of chromosomes, consistent with patterns found in other species [15], [16], [17]. To examine the recombination patterns in the regions with known introgressed segments from *M. Glaziovii*, we also constructed introgression dosage-specific genetic maps (Figure 1).

**Figure 2.**
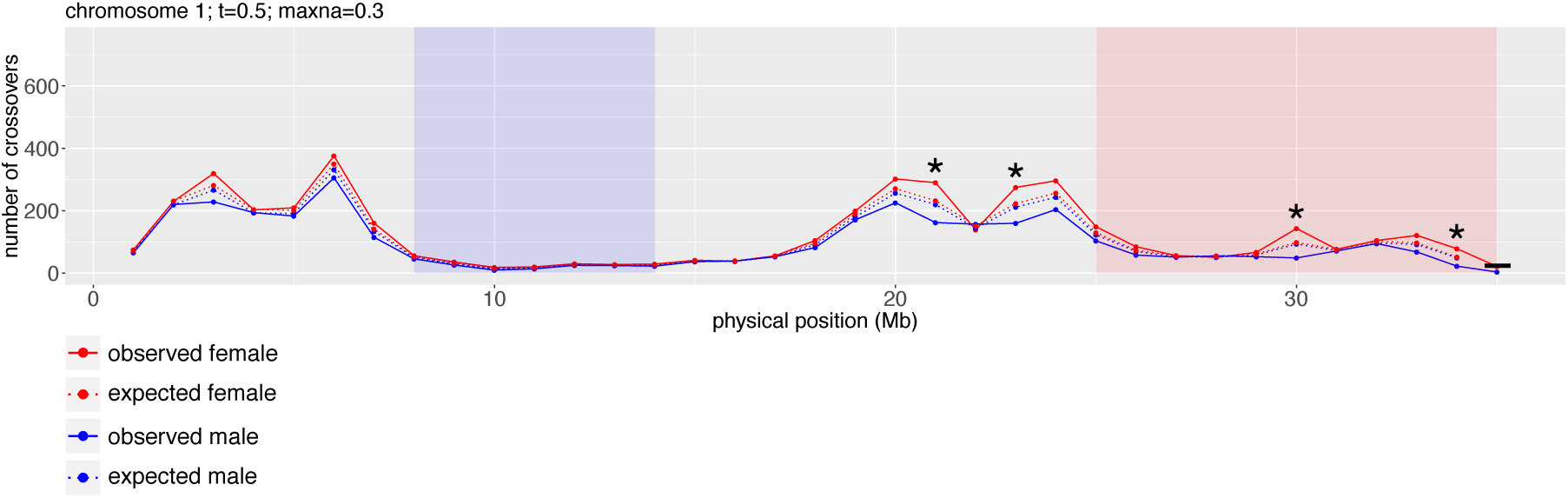
Distribution of crossover events across chromosome 1 for female meioses and male meioses. We divided each chromosome into 1-Mb windows and plotted the number of crossovers falling within each interval for female (red), and male (blue) meioses. Solid lines represent observed counts. Dashed lines represent expected counts, assuming that crossing over occurs with equal frequency in females and males. Asterisks show intervals with significantly different crossover frequency between male and female meioses. Dashes represent cases where we could not perform the chi-square test because the expected frequency count for one or more classes was less than five. We did not test for statistical significance in the last window of any chromosome since the last window is shorter than 1-Mb (no chromosome is perfectly divisible by 1-Mb). The centromere of the chromosome is shown in purple. The region with detected introgressions from *M. Glaziovii* is shown in red. We tested each interval at a significance level of *α*/*n*, where *α* = 0.05 and *n* = 506.

We also examined the crossover frequency in regions on chromosome 1 and 4 with known introgressions from *M. glaziovii*. For each region, we performed a chisquare test of equal counts to examine if individuals with zero, one, and two introgressed haplotypes experience different levels of recombination in the introgressed region. We performed the same test at the chromosome level to see if introgression class effects recombination levels across the entire chromosome. The chi-square test for introgressed regions on chromosomes 1 and 4 were significant with chromosome 1 having a p-value of 3.87e^-14^ and chromosome 4 having a p-value of 1.14e^-58^. Assuming that all individuals experience the same rate of recombination (regardless of introgression status), we expected to observe 536, 821, and 89 crossovers in individuals with zero, one, and two introgressions, respectively, but observed 680, 698, and 68 crossovers, respectively for chromosome 1. For chromosome 4, we expected to observe 1213 and 414 crossovers in individuals with zero and one introgression, respectively, but observed 1497 and 130, respectively. When performing the same test but on the non-introgressed portion of each chromosome, we found that for chromosome 4, individuals with different introgression statuses experienced significantly different levels of crossing over (p-value of 2.94e^-9^) but did not find this for chromosome 1 (p-value of 4.31e^-2^). These results indicate that the introgression on chromosome 4 affects recombination at the chromosomal level but the introgression on chromosome 1 affects only recombination in the introgressed region.

We tested if there is sexual-dimorphism in crossover number at the genome-wide level using a chi-square test of equal counts and found that the number of crossovers observed in male and female meioses significantly differed (p-value = 2.74 × 10^−8^). To investigate if crossover placement and rate varied between the two sexes in specific chromosomal regions, we examined the distribution of crossovers along each chromosome for female and male meioses separately. We divided each chromosome into windows of 1-Mb and plotted the number of crossovers detected in female meioses and male meioses in each 1-Mb window, with Figure 2 depicting that of chromosome 1 (Supplementary Figure 3 shows these plots for all 18 chromosomes).. Crossover placement along the chromosomes does not vary between male and female meiosis. To examine if crossover frequency in each window varied between the sexes, we performed a chi-square test of equal counts in each window. We did not test for statistical significance in the last window of any chromosome since the last window is shorter than 1-Mb. Of the 506 intervals tested, 45 (8.9%) passed the significance threshold. In these 45 intervals, female crossover count was significantly higher than expected and male crossover count was significantly lower than expected, a pattern observed in other taxa [18]. Statistically significant intervals did not consistently appear in any specific region of the chromosomes (Supplementary Fig 3).

We used the deviance goodness of fit test to test if crossover placements are random and independent events. The goodness of fit test was significant for all chromosomes except chromosomes 10, 17, and 18, indicating that the Poisson model does not fit the data observed on chromosomes 1-9 and 11-16 well and suggesting crossover interference.

## DISCUSSION

We used IITA’s multi-generational pedigree, consisting of 7,165 informative meioses (3,679 female; 3,486 male), to characterize recombination in cassava. Using SHAPEIT2 and duoHMM, we detected a total of 67,833 and 51,741 crossover-containing intervals from female and male meioses, respectively, across the 18 chromosomes. We used these crossovers to investigate (1) if the presence of wild introgressions from cassava’s wild relative *Manihot glaziovii* affects recombination rates in cassava and (2) if crossover number and spatial distribution differs between females and males.

In the 1930’s, breeders crossed cassava with its wild relative *M. glaziovii* in an effort to introduce cassava mosaic disease resistance into cassava. Wolfe et. al detected large *M. glaziovii* introgressions on chromosome 1, spanning from 25 Mb to the end of the chromosome, and on chromosome 4 from 5 Mb to 25 Mb [14]. We examined the recombination patterns in the introgression region on chromosome 1 and 4 by plotting separate genetic maps for individuals that are homozygous non-introgressed, individuals heterozygous for introgressions, and individuals that are homozygous introgressed in the regions (Figure 1). We observed that introgressions from *M. glaziovii* decreased recombination (Figure 1). We observed a dosage effect on chromosome 1, where having more introgression copies corresponded to lower recombination rates. This suggests the presence of a variant on the *M. glaziovii* haplotype that leads to lower overall recombination. For chromosome 4, none of our samples were homozygous introgressed, but we observed that heterozygous introgressed individuals experienced lower rates of recombination (the genetic map built from heterozygous introgressed individuals is shorter than the map built from homozygous non-introgressed individuals). Although individuals with one introgression undergo less recombination overall, we did observe that these individuals undergo more recombination leading up to the introgressed region (from 0 to 5 Mb) compared to individuals with zero introgressions. Recombination rate for these individuals then flattens close to zero for most of the introgressed region (recombination rate increases slightly in the region spanning 12 and 13 Mb and increases in the last 1 Mb of the region), whereas recombination continues to increase across the introgression region for homozygous non-introgressed individuals. Results from the chi-square test of equal counts for chromosome 1 and 4 showed that the level of crossing over between introgression classes is significantly different.

We observed similar spatial distributions of crossover events between the ICGMC map and our map, although it should be noted that we used a version of the ICGMC map as input when running SHAPEIT2 and duoHMM. Differences between our map and ICGMC’s could result from a number of reasons. The data used in our analysis was generated using a substantially different variant discovery pipeline than that used by ICGMC [19],[10], and the ICGMC map was generated using 10 nuclear families, each family consisting of 117 to 256 offspring. There is also the question of what value of *t* to use, as this dictates the number of crossovers available for map building. Using a higher *t* value results in more confident crossover intervals but also a lower number of crossovers for us to build a map with. We selected *t* = 0.5 because the developers of duoHMM found that at this threshold, the algorithm had a detection rate of 90.57% and a false discovery rate of 2.89% in simulations with realistic levels of genotyping error [11].

In this study, we used the multi-generational pedigree and GBS data from IITA to study recombination in cassava. We examined the recombination landscape across the 18 chromosomes of cassava and found that crossover rates vary greatly along the chromosomes and that all chromosomes except chromosomes 10, 17, and 18 displayed crossover interference. We constructed a genetic map using crossovers detected through SHAPEIT2 and duoHMM and compared it to ICGMC’s composite map. We also examined recombination in regions with known introgressed segments from cassava’s wild relative *M. glaziovii*. For both chromosome 1 and 4’s introgression region, we observed that individuals in different introgression classes experience significantly different rates of recombination within the introgressed region and that introgression-dosage affects recombination at the chromosome level for chromosome 4. Lastly, we looked at female and male meioses, separately and found that the spatial pattern of crossovers along the chromosomes did not vary between male and female meiosis at the qualitative level. We did, however, find evidence that female meioses undergo more recombination than male meioses, consistent with patterns found in other species [18].

## ACKNOWLEDGEMENTS

We acknowledge the Bill and Melinda Gates Foundation and the Department for International Development of the United Kingdom for funding this work through the “Next Generation Cassava Breeding Project”. We thank the entire Next Generation Cassava Breeding team for contributing to this study in the field and lab. We give special thanks to the International Institute of Tropical Agriculture (IITA), Ibadan, Nigeria for sharing their pedigree records with us for this study. Marnin Wolfe shared datasets on *M. glaziovii* introgressions with us, for which we are grateful.

